# Bilingualism Modulates Executive Function Development in Pre-School Aged Children: A Preliminary Study

**DOI:** 10.1101/2024.10.04.616691

**Authors:** Sally Sade, Scott Rathwell, Bryan Kolb, Claudia Gonzalez, Robbin Gibb

**Affiliations:** Canadian Centre for Behavioural Neuroscience, University of Lethbridge

## Abstract

This preliminary study was conducted to explore the effects of bilingualism on executive function development in children ages 3-5-years old. Two groups (bilinguals and monolinguals) were recruited across various sites in Southern Alberta. Children were assessed through parent rated executive function using the Behaviour Rating Inventory of Executive Function – Preschool version, a standardized assessment of executive function in children aged 2 years, 0 months through 5 years, 11 months. The questionnaire contains 63 items measuring 5 aspects of executive functioning, inhibit, shift, emotional control, working memory, and plan/organize. Children were also assessed using a battery of executive function tasks, which include the reverse categorization, pictorial Stroop, Dimensional Change Card Sort, backward digit span, and dyadic social play. Results show that bilingual children outperform monolinguals on the emotional control scale, dimensional change card sort and dyadic social play. Despite the controversial literature surrounding bilingualisms impact on executive function, the study reveals support for second language use to improve areas of executive function among young children.

## Introduction

One of the most profound, constant, and integrative human experiences modulating brain function is language use. For more than half of the world’s population, the intensive experience associated with language extends to managing two languages. In Canada specifically, 41.2% of individuals can communicate in more than one language (Statistics Canada, 2021). To date, research has shown that as bilingual individuals read, speak, or write in one language the second language (L2) is simultaneously active (see review in Dijkstra & van Heuven, 2002). Yet, bilingual individuals rarely report intrusion errors or confusion paralysis when trying to use a language. Instead, they are capable of inhibiting one language to successfully attend to another (Bialystok et al., 2009). Due to ongoing selection demands to inhibit the non-target language, this suggests attention networks undergo re-organization. The idea of bilingualism as a source of neuroplasticity builds on the adaptive control hypothesis which assumes there is an adaptation in domain general processes of attention subsequently shaping performance on nonverbal tasks recruiting such cognitive (executive) control (Green & Abutalebi, 2013). Decades of empirical evidence have established the positive effects of enriching experiences on rat behavior and learning (Hebb, 1949), however, the translatability to humans remained relatively unexplored until recent years. Some activities reported to induce changes in neural structures include playing the piano (Bengtsson et al., 2005), juggling (Draganski et al., 2004) and taxi-driving in London (Maguire et al., 2000). Thus, the adaptation of attentional control domain processes in bilinguals may elicit experience-dependent brain plasticity beyond language-specific processes.

Despite the prevalence of bilingualism, the research findings surrounding its cognitive impact on young children continues to be debated. Many early studies of child bilingualism led to the belief that L2 use was a social plague and detrimental to intelligence (review in Hakuta, 1986). One of these early studies, reported a negative relationship between the amount of foreign language used in homes and the median IQ of monolingual and bilingual school aged children (Goodenough, 1926). The study reported that the less foreign language children used (and the more English), the higher their IQ. The author interpreted this finding as L2 use being one of the key factors in producing mental retardation. The most influential of these early findings was a study comparing the vocabulary in a diverse linguistic population of 2- to 6-year-old children (Smith, 1939). The study reported that despite accounting for both languages, the total vocabulary of bilinguals was inferior to that of monolingual children. Later analyses point out several methodological flaws that applied to most early research on bilingualism: (a) group differences in age, socioeconomic status (SES) and sex were not accounted for, (b) the degree of bilingualism was determined by ‘foreignness’ of parents, (c) tests were solely administered in one language without acknowledging participants level of comprehension and (d) bilingualism was poorly defined and sometimes based on family names (Hakuta, 1986). Peal and Lambert (1962) shifted the perspective and introduced the notion that L2 use may positively impact cognitive performance in children. The study reported better performance by 10-year-old bilinguals on both verbal and nonverbal intelligence tests than their monolingual counterparts. The authors suggest performance differences due to bilingual children shifting between responses or tasks more efficiently compared to their monolingual counterparts. Although the study fell short to uneven socioeconomic status and the use of broadly based intelligence tests, it was the first study to provide evidence for a positive correlation between bilingualism in children and cognitive performance.

More recent studies have focused on the influence of bilingualism on higher-level cognitive processes known as executive function (EF), also called executive control or cognitive control. Various operational definitions of EF have been proposed, although none have been universally accepted. The common theme among these definitions is that EF is a construct describing multiple interrelated higher order regulatory cognitive processes required for self-regulation and goal-directed behavior. A prominent model of EF, known as the Unity and Diversity model, proposes that EF consists of discrete but interrelated “core” mental skills, including working memory -the ability to retain and manipulate information over a brief period of time for the purpose of making a response or completing a task; interference control – the ability to selectively attend to specific stimuli and focus attention for a prolonged period of time; inhibitory control - the ability to resist prepotent responses or distractions; and set shifting- the ability to swiftly adapt and adjust response sets as the situation demands (Miyake et al., 2000).

The framework suggests that higher order EFs such as planning, problem-solving and organization skills are built from these “core” executive processes. As this framework is the most cited, many studies utilize the Unity and Diversity model when defining EF. However, as research develops limitations of this model have been noted diminishing its validity (for more information see review – Sambal et al., 2023). More recent papers have considered other frameworks such as the Executive Control System (Anderson 2002, 2008), which conceptualizes EF as a system comprising four domains: attentional control, cognitive flexibility, goal setting, and information processing. Differences between the two frameworks are seen in the latter with the inclusion of a goal setting domain – the ability to initiate an activity or design an action plan to guide the completion of a task; and the information processing domain – which refers to the fluency, efficiency and speed of output. Lesion studies have linked the development of EF to the PFC. Furthermore, neuroimaging studies of healthy adults performing EF tasks report increased activation in the PFC, with some involvement of non-frontal regions. Activation of the non-frontal brain areas tend to be task dependent, whereas the prefrontal cortex serves as a major neural substrate of EF performance (Buchsbaum et al., 2005; Laird et al., 2005; Rottschy et al., 2012).

These skills begin to emerge during the perinatal period, strengthen significantly in early childhood-especially during the preschool years, and continue to refine through early adulthood (Shallice, 1988). Studies have shown that the level of EF development during the preschool years shape individual learning trajectories in school. Thus, a child’s “school readiness” and long-term academic or life success is highly correlated to EF performance during the preschool years. In fact, previous research has shown that EF is a better predictor of academic and life success than IQ (Alloway & Alloway, 2010; Willoughby et al., 2017). Although there is growing evidence that bilinguals perform better on nonverbal tasks measuring cognitive flexibility, inhibition and working memory across the lifespan, this stance remains controversial (see Bialystok, 2017; Pot et al., 2018 for reviews). The most recent meta-analysis assessed 152 studies to compare the relationship between bilingualism and EFs in adults (Lehtonen et al., 2018). Following correction for publication bias, the analysis found no bilingual advantage. Nichols and colleagues (2020) conducted a study on 11,041 participants measuring EF performance across a battery of EF tasks. The researchers stated that previous evidence lacked a sufficient sample size and was subject to publication bias. The study found no difference in EF between monolingual and bilinguals. Despite the large sample size, this study falls prey to many of the flaws of early research in this field. Firstly, the online survey simply asked participants whether they spoke a second language, without accounting for the age at which the L2 was acquired or the level of proficiency. Secondly, the study grouped an unusually wide age range into one category (i.e., 18-87). However, studies have shown that performance on EF tasks measuring inhibitory control through reaction time (RT) differ across age groups. More specifically, it has been shown that young adults (20-30 years old) with L2 exposure show no EF performance differences on the Simon Task, a measure of inhibitory control, whereas differences can be seen in children and older adults (Biaylstok et al., 2005). Thus, grouping large age ranges together filters out the possibility of finding significance. Lastly, the study did not include participants under the age of 18, overlooking a group of individuals who would be most impacted by experience-dependent plasticity.

Clarifying whether L2 exposure during early childhood is advantageous or detrimental is vital in establishing clear healthcare advice. Currently, the standard clinical advice for parents and/or caregivers is to abandon L2 use if the child is at high risk for developmental problems (Baralt & Mahoney, 2020). Research supporting this advice is limited and abstract. Thus, further evidence in this field is required to determine whether abandoning L2 use in high-risk populations is unwarranted. The aim of this preliminary study was to assess whether EF performance on a battery of tests in 3-5-year-old bilingual children differed than that of 3-5-year-old monolingual children.

## Methods Participants

Sixty-three children between the ages of 3 and 5 years old were recruited from 9 different sites (i.e., preschools and daycares) throughout southern Alberta. Of these, 6 participants were excluded due to missing measures or withdrawal from the study. Exclusion criteria for all participants included sensory impairments (i.e., blind or deaf) and inability to comprehend task instructions. After these exclusions, fifty-seven participants (34 females, 23 males) remained in the study (twenty-three 3-year-olds – *M* = 3.43, *SD* = ± .21; twenty-four 4-year-olds-*M*= 4.39, *SD* = ± .29; ten 5-year-olds-*M* = 5.37 *SD*= ± .35). In accordance with the Research Ethics Board at the University of Alberta (Study ID Pro00120179), written consent was obtained from the parents of the participants and verbal assent was obtained from the children prior to the start of the experiment.

## Procedure

Children were tested at their site on two occasions. Each testing session was performed less than 2 weeks apart, with each session lasting approximately 30-45 minutes. Testing sessions were divided into two rounds to ensure the child was not fatigued. Parents were asked to complete three paper-based questionnaires which include: (1) a participant information sheet used to collect demographic information including race, ethnicity, level of education, maternal, paternal and participants health (2) Behaviour Rating and Inventory Scale-Preschool Version (BRIEF-P) is a standardized rating scale of executive function for children between the ages of 2 years and 0 months to 5 years and 11 months. It includes 63 questions assessing a child’s behaviour over the past 6 months across five executive function clinical scales: Inhibit, Shift, Emotional Control, Working Memory, Plan/Organize (3) Language Experience and Proficiency Questionnaire which is a comprehensive report of the child’s exposure to all languages in the household and community completed by the parent. Considering the diverse range of languages (i.e., English, Swahili, Gujarti, Tagalog, French, Malayalaus, Spanish, Visayan, Portuguese, Karem, Russian, Ukrainian, Afrikaan, and Kalenjin) represented in the sample, the study focused on language exposure rather than language competence. The bilingual sample recruited were primarily simultaneous which is defined as L2 use starting from birth up to 18 months of age. As suggested by previous literature, children were categorized as bilingual if they receive at least 10-25% of exposure to L2 (Byers-Heinlein, place & hoff, 2011; Marchman et al., 2010; Marchman, Martinez-Sussmann & Dale, 2004). A subset of children had multiple language exposure to 3 (n = 4) languages. Since most children had exposure to 2 rather than 3 languages, the group is referred to as bilingual.

## Measures

### The Reverse Categorization Task

Block Sorting (adapted from Carlson et al., 2004) is a measure of cognitive flexibility and inhibitory control that consists of a pre-switch phase and a post-switch phase. The task consists of the following materials: a large bucket, a small bucket, 6 small blocks and 6 large blocks of the same colour (i.e., yellow). The experimenter places the large and small bucket in front of the child at a distance where they can reach but not look inside. In the pre-switch phase, the child was instructed to place the small block within the small bucket and the large block within the large bucket. The experimenter then began handing the blocks one-by-one using a pseudorandom order for a total of 12 trials. In the post-switch phase, the experimenter told the child they were going to play a silly version, where the large blocks would be placed into the small bucket and small blocks within the large bucket. The same procedure as the pre-switch phase was followed. For both phases, the child was given 1 practice round to ensure they understood the instructions. The child is scored based on the number of errors in the post-switch phase. For every error the child receives a score of 1 but if they correct themselves after making an error, they receive a score of 0.5. This conflict task is a measure of cognitive flexibility and inhibitory control as the child must inhibit using the previously learned rule and attend to a less apparent rule to perform the task correctly.

### Pictorial Stroop Task

The animal Stroop (adapted from Wright et al., 2003) is a measure of inhibitory control that consists of an identification phase and a Stroop phase. In the identification phase, the experimenter presented the child with an animal and asked them to identify the animal (3 x rabbit, 3 x cat, 3 x mouse, and 3 x horse) for a total of 12 trials presented in a pseudorandom order. In the Stroop phase, the experimenter told the child that they had some silly animals where the head and body of the animal differ. The experimenter presented the child with a mismatched body and head and asked them to identify the animal by naming solely the body. The child’s understanding of the rules was confirmed through 4 practice trials prior to proceeding. The practice trials included (head-body): rabbit – horse, cat– mouse, horse-rabbit, and mouse-horse allowing the child to inhibit naming the head for each animal. Mismatched body and heads were randomly selected and presented to the child in a pseudorandom order for a total of 12 trials. The child is scored based on the number of errors in the Stroop phase. For every error the child receives a score of 1 but if they correct themselves after making an error, they receive a score of 0.5. This conflict task is a measure of inhibitory control because the child needs to inhibit the distracting head and focus on the relevant body to engage in the task correctly.

### Grass and Snow

The Grass-Snow task is a Stroop-like task (adapted from Gerstadt et al., 1996) a measure of inhibitory control that consists of an identification phase and a Stroop phase. Materials required include a green and white laminated 8.5 × 11-inch sheet of paper. In the identification phase, both sheets of paper were placed in front of the child, and they were asked to identify which paper resembled “grass” and which resembled “snow.” The children were instructed to point to the white paper when the experimenter said “snow” and point to the green paper when they said “grass” for 10 trials (5 x grass, 5 x snow). The experimenter read a pseudorandom list. In the Stroop phase, the experimenter told the child they were going to play a silly version where the white piece of paper represents “grass”, and the green piece of paper represents “snow”. The child was given a practice trial to ensure they understood the rules prior to beginning. The experimenter followed the same procedure for a total of 10 trials. The child is scored based on the number of errors in the Stroop phase. For every error the child receives a score of 1 but if they correct themselves after making an error, they receive a score of 0.5. This conflict task requires the child to inhibit the urge to point at the color matching the word and instead point to the incongruent sheet.

### Dimensional Change Card Sort (DCCS)

Turtles and Trains (adapted from Zelazo, 2006) is a variation of the Dimensional Change Card Sort (DCCS), a standard procedure for measuring executive function in children. The task consists of a standard and challenging border version. In the standard task, participants sort bivalent cards (i.e., purple train or orange turtle) by one dimension (e.g., colour) for 6 trials and then are instructed to switch and sort the same cards in a new dimension (e.g., shape) for a total of 12 trials. If the child successfully completed the standard version the experimenter proceeded to a more challenging border version. In the border version, participants are instructed to sort bivalent cards with a black 3-mm border (n = 6) according to the shape dimension and bivalent cards without a border (n = 6) according to the colour dimension for a total of 12 trials. Prior to the border version the participant had 4 practice trials in which corrective feedback was provided if needed. Children were scored based on the number of errors. This task measures cognitive flexibility, working memory and inhibitory control because the child is required to inhibit conflicting dimensions, shift between the previously learned rules and retain and update the new rule to engage in the task correctly.

### Backward Digit Span

The backward digit span (adapted from Wechsler, 2008 & Shen et al., 2020) is a measure of working memory or updating. Participants are given a practice trial using the numbers 1,2 with corrective feedback. The experimenter began the task, once the participant had successfully mastered the concept of repeating a sequence of numbers in the reverse order. The experimenter reads aloud a sequence of two digits and asks the child to repeat the sequence in reverse. If successful, the experimenter continues to read aloud sequences in increasing length (up to a maximum of 4 digits). If the participant was unsuccessful after two repetitions the task is ended. All participants were asked to repeat the same sequence of numbers that had been randomly selected from 0-9. Children were scored in accordance with Shen et al. (2020). This task is an indicator of working memory or updating as the participant is required to retain and manipulate the series of digits to complete the task correctly.

### Dyadic Social Play

The dyadic unstructured social play task is a measure of the participants ability to navigate social interactions (Yusuke, 2014). Participants are randomly paired from the same site and presented with a plethora of large blocks in which they are encouraged to construct on object of their choice while working together for 5 minutes. Participants are scored on a scale (adapted from Howes and Matheson’s Peer Play Scale) from 0-4 in three different categories (type of play, type of interaction and type of engagement). Type of play refers to the participants ability engage in an “enriched” play filled with collaboration and negotiation. Type of interaction refers to the participants ability to focus on the partner as the primary focus of the play rather than the blocks. Type of affects refers to the participants overall emotional engagement with the other youth which include eye contact, smiling and some apparent enjoyment. Participants may receive a maximum score of 4 for each category. This task assesses the participants level of social interaction which can influence EF skills in young children (Yusuke, 2014).

## Results

### Behaviour Rating Inventory of Executive Function

BRIEF-P scores were calculated using the procedure provided in the BRIEF-P manual (Gioia et al., 2003). A raw score was obtained and converted into a t-score, where lower t-scores are associated with better EF performance across all subscales (inhibit, shift, emotional control, working memory, and plan/organize, Inhibitory Self-Control Index (ISCI), Flexibility Index (FI), Emergent Metacognition Index (EMI) and Global Executive Composite (GEC). For this task, a non-parametric Mann Whitney U test was appropriate to use, given BRIEF-P scores are standardized which adjusts for across age cohort comparison, no within-subject variables, and the non-normal distribution across all variables of the data. Descriptive statistics of BRIEF-P scores across language groups can be found below in Table 1.

**Table 1.** Descriptive statistics of parent-rated EF across language groups.

| BRIEF-P | Monolinguals |  |  | Bilinguals |  |  |
| --- | --- | --- | --- | --- | --- | --- |
| | | $n = 30$ | | | $n = 18$ | |
|  | *M* | *SD* | *Md* | *M* | *SD* | *Md* |
| Inhibit | 50.97 | 10.82 | 50.00 | 46.61 | 7.42 | 46.50 |
| Shift | 51.17 | 12.35 | 49.00 | 45.44 | 5.85 | 46.00 |
| Emotional Control | 52.97 | 10.11 | 52.50 | 44.61 | 7.37 | 44.50 |
| Working Memory | 52.33 | 12.44 | 50.50 | 49.28 | 10.16 | 46.00 |
| Planning/Organization | 51.57 | 13.62 | 51.00 | 46.17 | 6.90 | 45.50 |
| ISCI | 51.93 | 10.69 | 52.00 | 45.50 | 7.24 | 45.50 |
| FI | 52.33 | 11.23 | 52.00 | 44.00 | 7.16 | 41.00 |
| EMI | 52.23 | 13.60 | 50.50 | 46.94 | 8.31 | 45.00 |
| GEC | 51.80 | 12.47 | 49.00 | 45.39 | 7.83 | 43.50 |

These analyses were performed on SPSS version 28, and eta squared was calculated to estimate the effect size (η^2^ = *Z* ^2^ ÷ *N*-1). To interpret effect size magnitude, we followed Cohen’s guidelines where η^2^ = .01 is considered small, η^2^ = .06 is medium and η^2^ = .14 is large.

A Mann-Whitney U test revealed that BRIEF-P scores varied across subcomponents of EF. Emotional Control scores were significantly lower in the bilingual group (*Md* = 44.50, *n* =18) compared to the monolingual group (*Md* = 52.50, *n* = 30), *U* = 130.50, *Z*= -2.977, *p* = .003 with a medium effect size η^2^ = 0.13 ; Inhibitory Self-Control Index were significantly lower in the bilingual group (*Md* = 45.50, *n* = 18) compared to the monolingual group (*Md* = 52.00, *n* = 30), *U* = 169.50, *Z*= -2.176, *p* = .030 with a medium effect size η^2^ = .09; and Flexibility Index were significantly lower in the bilingual group (*Md* = 41.00, *n* = 18) compared to the monolingual group (*Md* = 52.00, *n* = 30), *U* = 143.50, *Z*= -2.698, *p* = .007 with a medium effect size η^2^ = 0.12. As shown in Table 2, no significant differences were found across other BRIEF-P subcomponents.

**Table 2.** Language group comparisons on parent-rated EF.

| BRIEF-P | Mann-Whitney U | Z | Asymp. Sig (2-tailed) |
| --- | --- | --- | --- |
| Inhibit | 208.500 | -1.312 | .190 |
| Shift | 201.500 | -1.461 | .144 |
| Emotional Control | 130.500 | -2.977 | **.003** |
| Working Memory | 222.000 | -.896 | .370 |
| Planning/Organization | 196.500 | -1.572 | .116 |
| ISCI | 168.500 | -2.176 | **.030** |
| FI | 143.500 | -2.698 | **.007** |
| EMI | 207.000 | -1.343 | .179 |
| GEC | 183.000 | -1.854 | .064 |

### Dyadic Social Play

The data obtained from the dyadic social play across groups satisfied the assumptions of a parametric test including normality, homogeneity of variance and independence. Thus, a two-way ANOVA was performed to evaluate the effects of language group and age on social play score. The means and standard deviations for social play score are presented in Table 3 below. Estimated marginal means of social score across language group and age are presented in Figure 1.

**Table 3.** Descriptive statistics for social play score.

| Language Group | Age | *M* | *SD* | N |
| --- | --- | --- | --- | --- |
| Monolingual | 3.00 | 5.7222 | 1.85592 | 9 |
|  | 4.00 | 6.1429 | 1.63411 | 14 |
|  | 5.00 | 6.3000 | 1.44049 | 5 |
|  | Total | 6.0357 | 1.63259 | 28 |
| Bilingual | 3.00 | 5.0500 | 1.32183 | 10 |
|  | 4.00 | 8.5625 | 1.87916 | 8 |
|  | 5.00 | 8.6500 | 2.14243 | 4 |
|  | Total | 6.9818 | 2.41673 | 22 |
| Total | 3.00 | 5.3684 | 1.58852 | 19 |
|  | 4.00 | 7.0227 | 2.06142 | 22 |
|  | 5.00 | 7.3444 | 2.07190 | 9 |
|  | Total | 6.4520 | 2.04862 | 50 |

**Figure 1.**
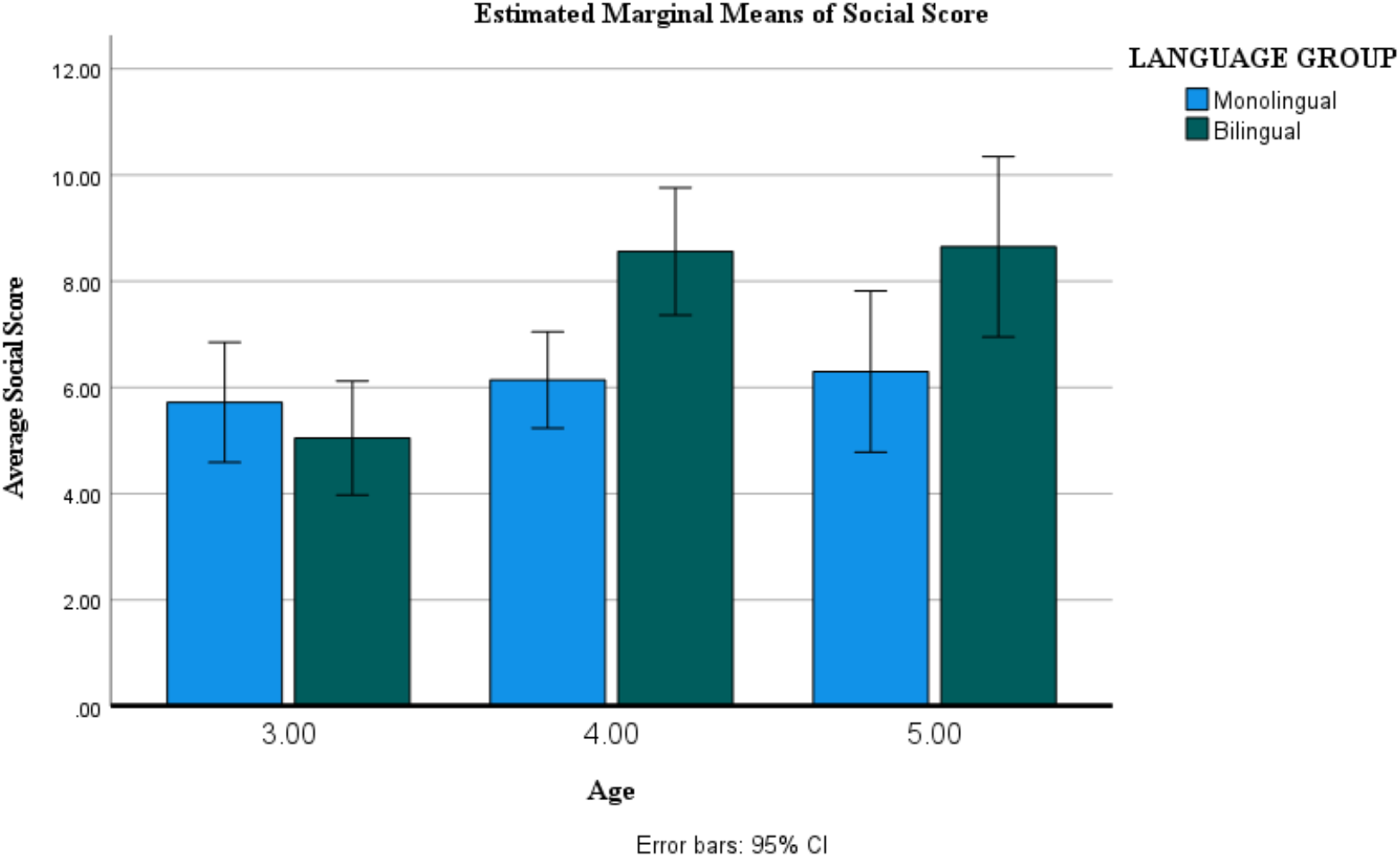
Mean performance on social play task across ages by language group

The results indicate a significant main effect for language group, *F*([1], [44]) = [6.903], p [.012], partial η^2^ = [.136]; a significant main effect for age, *F*([2], [44]) = [8.16], p [<.001], partial η^2^ = [.270]; and a significant interaction between language group and age, *F*([2], [44]) = [4.78], p [.013], partial η^2^ = [.179].

Post hoc testing using least significant difference (LSD) indicated that social play score was significantly higher for 4-year-old bilinguals than 3-year-old bilinguals (p= [<.001]; and significantly higher for 5-year-old bilinguals than 3-year-old bilinguals (p= [<.001]). There was no significant difference between social play score across 3-year-old monolinguals and 4-year-old monolinguals (p= [.562]); and no significant difference between 3-year-old monolingual children and 5-year-old monolingual children (p = [.542]).

### Battery of EF Tests

Distributions for each EF task did not satisfy the assumptions of a parametric test. Data were positively skewed and did not satisfy homogeneity of variance. Thus, a Mann – U Whitney test comparing effects between language group and EF performance was computed for each age group. Analysis was conducted separately between age groups as the table-top assessments are unable to correct for age. The study reports significant findings obtained from the Dimensional Change Card Sort standard version. No significance was found across age groups for the Backward Digit Span, Grass and Snow, Animal Stroop, Block Sorting Tasks, Dimensional Card Sort-Border Version. The non-significant result could be due to low statistical power, as shown in Table 4 sample size varied across ages which may increase the impact from outliers.

**Table 4.** Language group comparisons across age group of Dimensional Change Card Sort: Switch Percent Errors.

| Age | N | Mann-Whitney U | Z | Asymp. Sig (2-tailed) |
| --- | --- | --- | --- | --- |
| 3.00 | 18 | 44.00 | -1.27 | .205 |
| 4.00 | 20 | 28.00 | -2.312 | **.021** |
| 5.00 | 9 | 7.50 | -.732 | .464 |

The Dimensional Change Card Sort: Standard version percent errors were significantly lower in the 4-year-old bilingual group (*Md* = .00, *n* =8) compared to the 4-year-old monolingual group (*Md* = 41.67, *n* = 14), *U* = 28.00, *z*= -2.312, *p* = .021 with a large effect size η^2^ = 0.22. As shown in Table 4, these results were not found among the 3-year-old and 5-year-old age group.

## Discussion

This preliminary study sought to examine whether 3-5-year-old bilingual children perform better on EF tasks compared to monolingual children. Data from this study showed that the bilingual children scored significantly better than their monolingual peers on parent rated BRIEF-P scales of emotional control, inhibitory self-control index and flexibility index. The emotional control scale measures a child’s ability to modulate emotional responses. For instance, children capable of not having an outburst to seemingly minor events can be perceived as having good emotional control. The inhibitory self-control index is a combination of emotional control and inhibition. It represents a child’s ability to appropriately self-regulate. The flexibility index is a combination of shifting and emotional control. It represents a child’s ability to shift between actions, emotions and behavior. Furthermore, bilingual children scored significantly better than their monolingual counterparts on the dyadic social play task. The magnitude of these effects significantly increased across 3- and 5-year-old bilinguals compared to monolinguals. This suggests that as bilingual children age their social skills increase at a higher rate than that of their monolingual counterparts. These results align with previous studies assessing social play and language abilities (Holmes et al., 2014). However, this is the first study to show a difference among dyadic social play in bilingual and monolingual children. Lastly, 4-year-old bilingual children performed better on the standard version of the Dimensional Change Card Sort task compared to their monolingual peers. No differences were found amongst other table-top assessments of EF.

The present study conflicts with the results of Bialystok (2015), who found better performance of bilinguals on tasks requiring inhibitory control. One possible explanation for the difference in results could be that the present study lacks sufficient power due to variations in sample size across ages. Although EF is measured as a unified construct, the task impurity problem posits that not all tasks will measure EF skills equally. Previous studies report better inhibitory control in bilinguals on the Simon and Flanker task, however, this study used alternative tasks to measure inhibitory control. Given that the adaptive control hypothesis suggests an adaption in domain general processes of attentional control, further research should address why bilinguals do not perform better in comparison to monolinguals on all non-verbal tasks measuring such executive controls.

In addition, the present study assessed 3-5-year old bilingual children with 10-25% of L2 use prior to 18 months. The controversial findings surrounding cognitive improvements and bilingualism may stem from the lack of an operational definition for bilingualism. The arbitrary definition of 10-25% may not accurately depict the variety in which L2 use exists in each individual. Alternatively, some studies have defined bilingualism as a categorical variable (i.e. yes or no). Changing the perspective to assessing bilingualism on a continuous scale rather than categorical through gathering a comprehensive understanding of the child’s language background. Factors that may influence bilingualism include (1) how often they speak or hear the second language at home, (2) how often they speak or hear the second language at school, (3) how often they speak or hear the second language in the community, (4) is the second language heavily influenced through family or the community, (5) on a percentage scale how often does each parent speak a particular language to their child. Future studies consider calculating a ratio of first and second language use depending on the factors stated above.

## Notes

### Competing Interest Statement

The authors have declared no competing interest.

